# Sperm induction of somatic cell-cell fusion as a novel functional test

**DOI:** 10.1101/2023.07.18.549574

**Authors:** Nicolas G. Brukman, Clari Valansi, Benjamin Podbilewicz

## Abstract

The fusion of mammalian gametes requires the interaction between IZUMO1 on the sperm and JUNO on the oocyte. We have recently shown that ectopic expression of mouse IZUMO1 induces cell-cell fusion and that sperm can fuse to fibroblasts expressing JUNO. Here, we found that the incubation of mouse sperm with hamster fibroblasts or human epithelial cells in culture induces the fusion between these somatic cells and formation of syncytia, a pattern previously observed with some animal viruses. This sperm-induced cell-cell fusion requires a species-matching JUNO on both fusing cells, can be blocked by an antibody against IZUMO1, and does not rely on the synthesis of new proteins. The fusion is dependent on the sperm’s fusogenic capacity, making this a reliable, fast and simple method for predicting sperm function during the diagnosis of male infertility.

**Highlights:**

- Sperm induces viral-like fusion of somatic cells expressing JUNO
- We developed a new technique to determine the fertilization potential of sperm
- The test measures the capacity of sperm to induce somatic cell fusion
- The degree of somatic cell fusion correlates with the fertilizing ability of sperm

**Graphical abstract:** 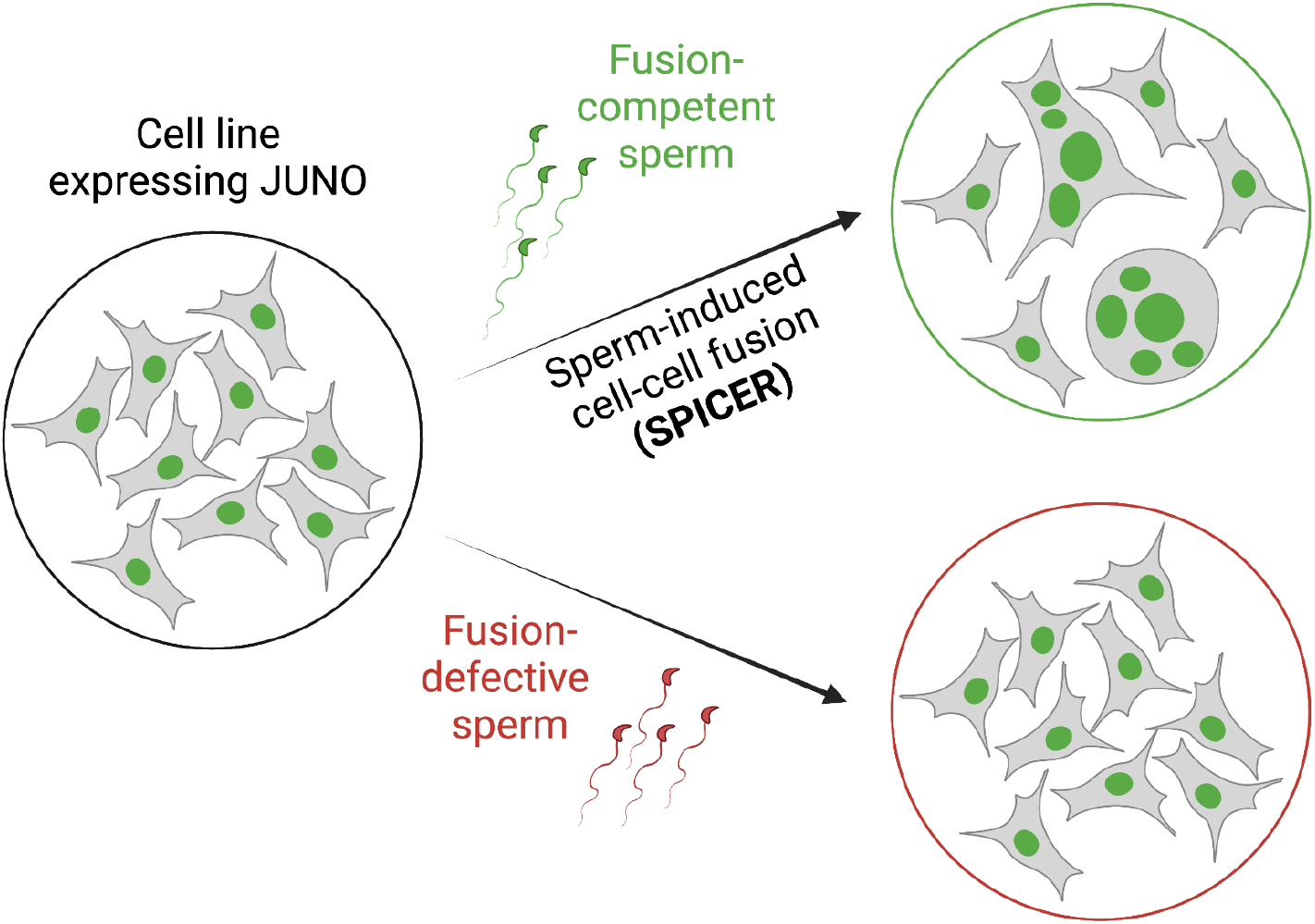

## Introduction

Infertility is estimated to affect approximately 15% of the population (World Health Organization 2023), and many couples dealing with this burden turn to assisted reproductive techniques (ARTs) as a possible solution (Steptoe and Edwards 1978; Palermo et al. 1992). During fertility evaluation of a male patient, a basic semen examination is performed, where different sperm parameters are determined: concentration, motility, morphology and vitality (World Health Organization, HRP 2021). This basic analysis can be complemented with additional tests to determine the fertilizing potential of the sperm and choose the best treatment for the couple. The hamster oocyte penetration (HOP) test has been proposed as a quantitative method for analyzing the fusogenic potential of human spermatozoa (Aitken and Elton 1986; R. Yanagimachi, Yanagimachi, and Rogers 1976; Zainul Rashid et al. 1998), however, it was excluded from the newest WHO laboratory manual for the examination and processing of human semen (World Health Organization, HRP 2021) for being considered obsolete. Therefore, to date there is no other standardized methodology to analyze specifically the ability of sperm to fuse to oocytes.

In mammals, the adhesion of the sperm to the oocyte plasma membranes is mediated by the species-specific interaction of two membrane proteins: IZUMO1 and JUNO (Bianchi and Wright 2015; Aydin et al. 2016; Ohto et al. 2016; Matsumura et al. 2021). The transmembrane protein IZUMO1 is expressed during spermatogenesis and localizes to the fusogenic region of the sperm head after an exocytic process named acrosome reaction (Inoue et al. 2005; Satouh et al. 2012; Inoue et al. 2015; K. Nishimura et al. 2016). On the other hand, the IZUMO1 receptor, JUNO, is an oocyte protein bound to the plasma membrane by a GPI lipid anchor (Bianchi et al. 2014). The subsequent fusion of the two gametes relies on the action of IZUMO1 in a unilateral manner (Brukman et al. 2023). The IZUMO1-JUNO interaction was characterized also in humans (Aydin et al. 2016; Ohto et al. 2016) and its relevance for human fertility is supported by the presence of antibodies against IZUMO1 in the sera of immuno-infertile women ((Clark and Naz 2013; Yu et al. 2018; Enoiu et al. 2022)), the mutations on JUNO in cases of fertilization failure and polyspermy ((Clark and Naz 2013; Yu et al. 2018; Enoiu et al. 2022)), and the lower levels of IZUMO1 observed in sperm that failed to fertilize during *in vitro* fertilization (IVF) treatments (Clark and Naz 2013; Yu et al. 2018; Enoiu et al. 2022).

Recently, we found that mouse sperm can fuse to fibroblasts ectopically expressing the sperm-receptor JUNO (Brukman et al. 2023). Here, we show that after sperm fuses with a somatic cell, this cell can fuse with additional cells inducing syncytia formation - a single cell with several nuclei (multinucleated cell). This is possibly mediated by the bridging of a single sperm simultaneously fused to two different cells. We call this process “SPerm-Induced CEll-cell fusion Requiring JUNO” (SPICER). We found that sperm with higher fertilizing ability can induce the fusion of somatic cells more efficiently, judged by the increased levels of multinucleation. This establishes the basis for the future development of a new method for diagnosis of male fertility centered on the ability of sperm to induce cell-cell fusion *in vitro*.

## Results

### Sperm fusion to fibroblasts promotes syncytia formation

There are reports showing that mammalian sperm can fuse to somatic cells (Rival et al. 2019; Mattioli et al. 2009; Bendich, Borenfreund, and Sternberg 1974). In our experimental conditions, mouse sperm cells only fuse to Baby Hamster Kidney (BHK) cells if they are induced to express the sperm-receptor, JUNO (Brukman et al. 2023). This fusion was demonstrated by detecting the transfer of the DNA binding GFP (GFP-MBD) from the BHK cells to the sperm heads (Brukman et al. 2023). To further study the mechanisms of mammalian sperm-oocyte fusion, we incubated mouse sperm with BHK cells expressing JUNO (functioning as pseudo-oocytes) and determined the efficiency of the sperm-BHK cell interactions. Surprisingly, we found that sperm induces the formation of multinucleated BHK cells (syncytia; Fig. 1A-C). The induction of multinucleation was dependent on the presence of JUNO (Fig. 1B-C), however, JUNO expression alone is not sufficient to induce this process. Only cells with sperm fused to them form syncytia (Fig. 1C). Furthermore, larger syncytia tend to contain more sperm (Fig. S1A) and the levels of multinucleation were dependent on the amount of sperm added to the cells (Fig. S1B). Thus, the fusion of sperm with JUNO-expressing BHK cells is required for inducing subsequent multinucleation of these somatic cells. This phenomenon was unexpected, leading us to further study the mechanisms of this process.

**Figure 1:**
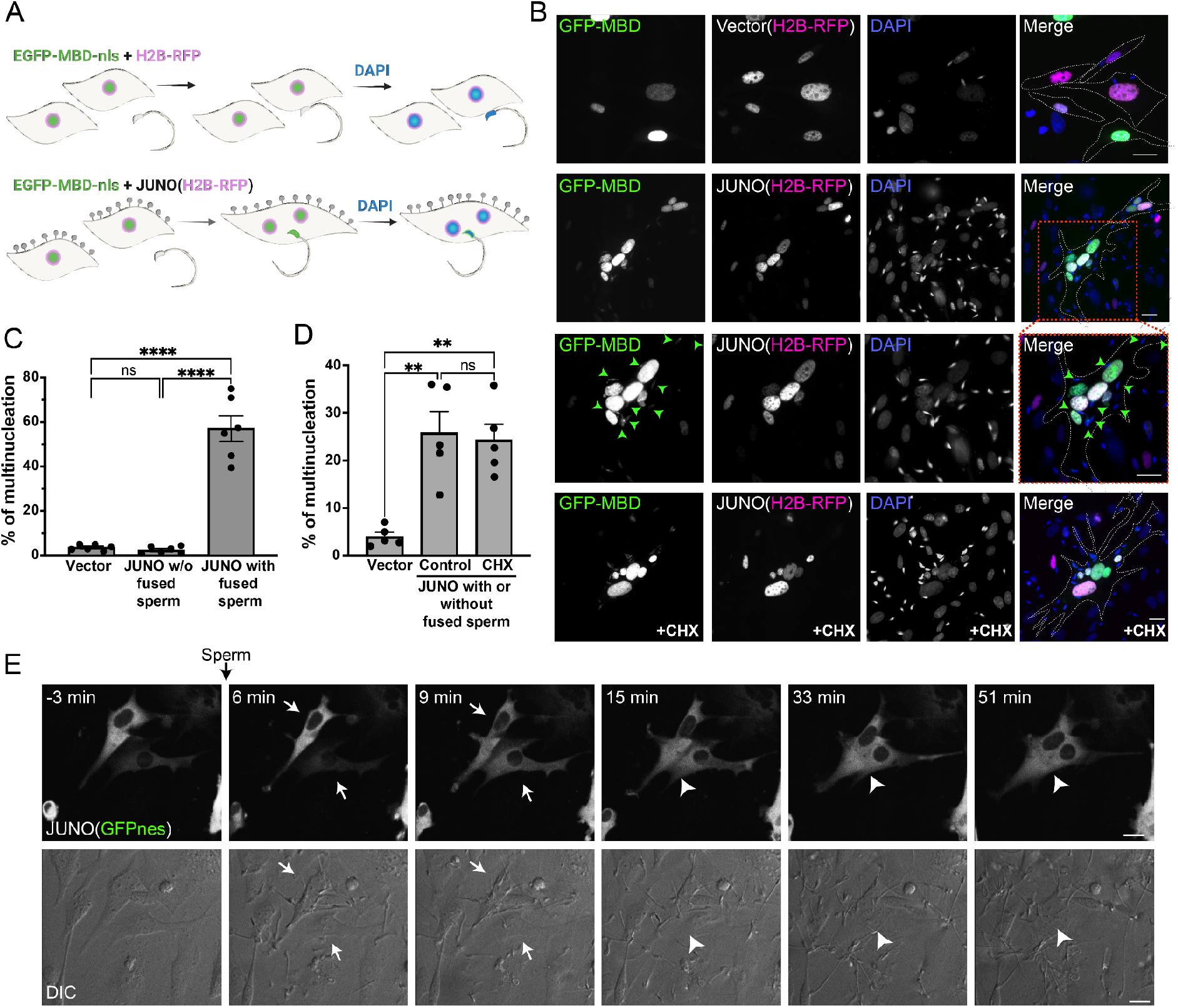
Sperm induce syncytia formation of fibroblasts. **(A)** Scheme of experimental design: Baby Hamster Kidney (BHK) cells were transfected with either pCI::H2B-RFP or pCI::JUNO::H2B-RFP vectors and pcDNA3.1-EGFP-MBD-nls. Mouse sperm were obtained from the epididymis of adult mice and capacitated in HTF capacitating medium. Sperm cells were co-incubated with the BHK cells for 4 h and then the cells were fixed and stained with DAPI to detect the DNA. Grey lollipops represent JUNO molecules. **(B)** Representative images showing H2B-RFP (magenta), GFP-MBD (green), DAPI (blue) channels and the merge. Dotted lines contour relevant cells. Green arrowheads point to fused sperm (GFP positive). The presence of 200 µg/ml of cycloheximide is indicated (CHX). Scale bars, 20 µm. **(C)** The percentage of multinucleation was defined as the ratio between the nuclei in multinucleated cells (NuM) and the total number in nuclei fluorescent cells (NuF), as follows: % of multinucleation = (NuM/NuF) × 100. We show individual data and means ± SEM of six independent experiments. The number of nuclei counted per experiment and per treatment was 500. For JUNO transfected cells, multinucleation was counted separately for cells with and without sperm fused to them. Comparisons were made with one-way ANOVA followed by Tukey’s test. ns = non-significant, ****P < 0.0001. **(D)** In another set of experiments JUNO-transfected cells were treated with 200 µg/ml CXH to inhibit *de novo* synthesis of proteins. Multinucleation was quantified for the whole population of transfected cells. We show individual data and means ± SEM of five independent experiments. ns = non-significant, **P < 0.01. **(E)** Time-lapse images from a movie showing sperm-induced cell-cell fusion. BHK cells were transfected with the pCI::JUNO::GFPnes plasmid and sperm were added at time=0 min. Arrows and arrowheads indicate contacting and fused cells, respectively. The green channel (GFPnes) and the DIC images are shown (see also Movie S1). Scale bars, 20 µm.

### Sperm induce syncytia formation using a viral-like mechanism

To our knowledge, sperm-induced cell-cell fusion has not been described in any species. However, decades ago, it was first described that some somatic cells can fuse following viral infection (Okada 1962; Kohn 1965) and later confirmed for diverse viruses, including SARS-CoV2 (Buchrieser et al. 2020). This process may require the synthesis of new viral proteins and therefore induces Fusion From Within (FFWI) while other viruses induce fusion independently of protein synthesis in a process called Fusion From Without (FFWO) (Bratt and Gallaher 1969; Duelli and Lazebnik 2007). For instance, orthoreoviruses induce the expression of fusion-associated small transmembrane (FAST) proteins upon infection, promoting FFWI (Ciechonska, Key, and Duncan 2014; Theuerkauf et al. 2021). We hypothesized that in a way reminiscent of viral induced cell-cell fusion, sperm could induce syncytia formation following its merger via FFWI or FFWO. To distinguish between these alternative mechanisms, we performed the experiment in the presence of cycloheximide to inhibit protein synthesis. We found no differences in the levels of multinucleation between control and cycloheximide treatments (Fig. 1B and D). This shows that *de novo* protein synthesis is not required for the induction of multinucleation by sperm and suggests a mechanism of FFWO. Additionally, live imaging of the process showed that fusion between cells occurs efficiently within 10-15 minutes after the addition of the sperm (Fig. 1E, Movie S1), suggesting that protein synthesis is not involved. Even though it is not known whether sperm can induce fusion of somatic cells *in vivo*, we show that sperm use a viral-like mechanism of FFWO when the fibroblasts express the oocyte JUNO *in vitro*.

### Syncytia formation results from the fusion of cells expressing JUNO

Then we asked whether the sperm-induced multinucleation was a consequence of BHK-BHK fusion and if so, whether it requires JUNO to be present on both fusing cells. For that, we employed a content mixing experiment where two populations of cells expressing different fluorescent markers were mixed and then exposed to sperm (Fig. 2A). We observed the formation of multinucleated hybrid cells containing both fluorescent markers, confirming the fusion of BHK cells expressing JUNO (Fig. 2B). This BHK-BHK fusion was not observed when only one or neither of the cell populations expressed JUNO (Fig. 2B-C), indicating that sperm-induced multinucleation is indeed BHK-BHK fusion which relies on bilateral expression of JUNO. An alternative explanation for sperm-induced multinucleation is a failure in cytokinesis. However, the presence of an inhibitor of the cell cycle (FdUdr, 5-fluoro-2′-deoxyuridine, (Valansi et al. 2017)) does not inhibit multinucleation (Fig. 3A-B), ruling out that syncytia formation is as a consequence of failure in the division of the BHKs and confirming the occurrence of fusion between them. Following these findings, we named this process SPICER that stands for “SPerm-Induced CEll-cell fusion Requiring JUNO”.

**Figure 2:**
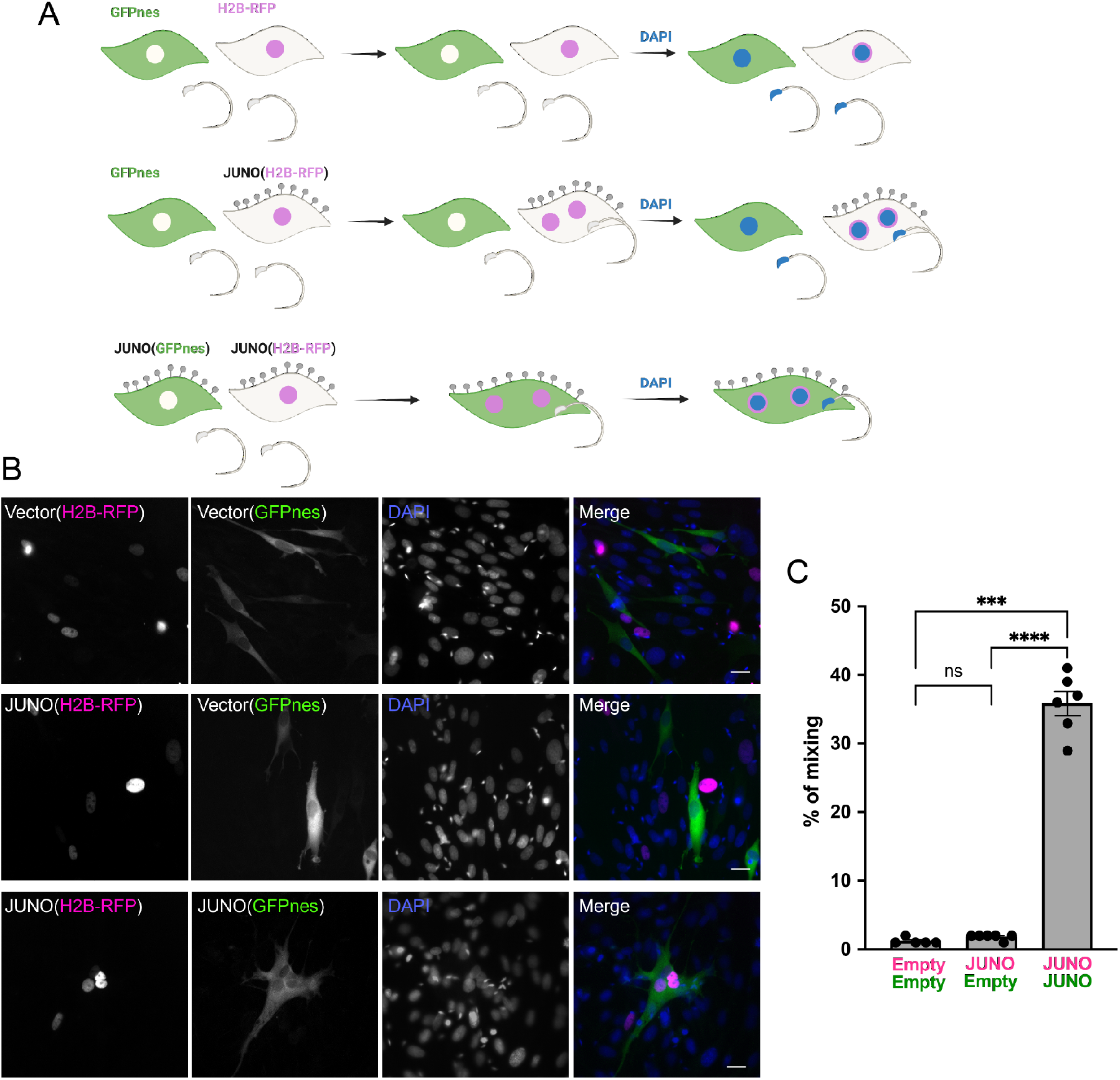
Sperm-induced fusion is dependent on JUNO. (**A**) Scheme of experimental design. BHK cells transfected with pCI::GFPnes or pCI::H2B-RFP empty vectors or containing the coding sequence for the expression of JUNO were mixed as indicated. Later, the cells were co-incubated with sperm for 4 h, fixed and stained with DAPI. (**B**) Representative images for each treatment. Each separate channel for GFPnes (green cytoplasm), H2B-RFP (magenta nuclei) and DAPI staining (blue) are shown. Scale bars, 20 µm. Fused, mixed cells contain both GFPnes and H2B-RFP staining. (**C**) Quantification of content mixing experiments. The percentage of mixing was defined as the ratio between the nuclei in mixed cells (NuM) and the total number of nuclei in mixed cells and fluorescent cells whose cell bodies were in contact that did not fuse (NuC), as follows: % of mixing = (NuM/[NuM + NuC]) × 100. Bar chart showing individual experiment values (each corresponding to 1,000 nuclei) and means ± SEM of six independent experiments. Comparisons by one-way ANOVA followed by Tukey’s test. ns = non-significant, ***P < 0.001, ****P < 0.0001.

**Figure 3:**
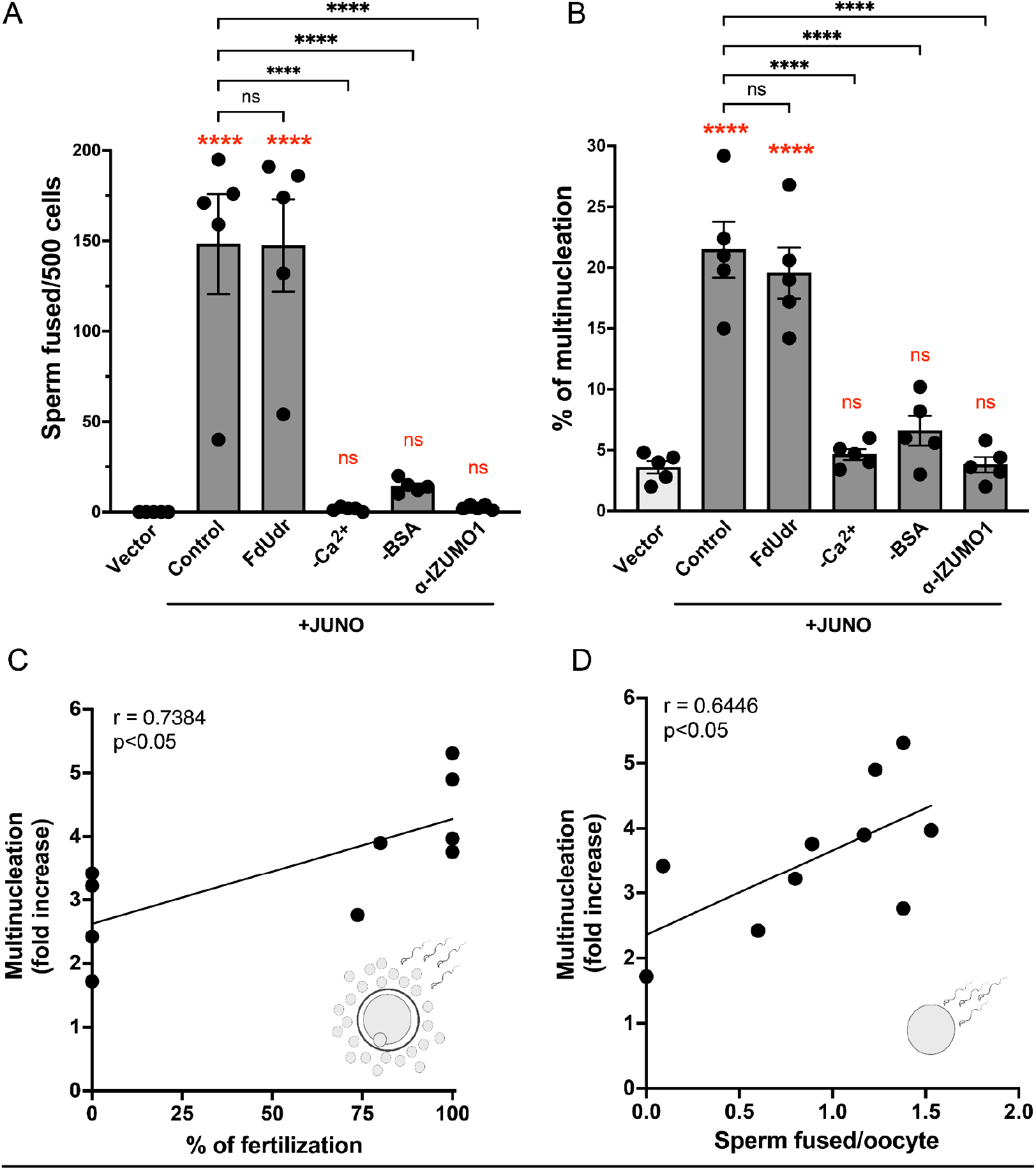
Syncytia formation requires a functional sperm, is dependent on IZUMO1 and is not affected by an inhibitor of cell division. BHK cells were transfected with pcDNA3.1-EGFP-MBD-nls together with pCI::H2B-RFP (empty vector) or pCI::JUNO::H2B-RFP and co-incubated with control sperm in the presence of 20 µM of the inhibitor of cell division FdUdr or of 1 µg/µl of anti-IZUMO1 antibody (clone Mab120). Alternatively, cells were mixed with sperm incubated in a medium lacking calcium (-Ca^2+^) or bovine serum albumin (-BSA); both conditions fail to support sperm capacitation. The number of sperm fused per 500 fluorescent cells **(A)** and the percentage of multinucleation **(B)** were determined. Bar charts showing individual experiment values and means ± SEM of five independent experiments. Comparisons by one-way ANOVA followed by Tukey’s test. In red are the comparisons against the empty vector. ns = non-significant, ***P < 0.001, ****P < 0.0001. **(C-D)** Multinucleation levels relative to the control without sperm as a function of the percentage of fertilized eggs when cumulus-oocytes complexes were used (**C**), or as a function of the number of sperm fused per oocytes when ZP-free eggs were employed (**D**). Each dot corresponds to a different mouse. The Pearson’s coefficient ‘r’ and the significance are included in each panel.

### Sperm fuses simultaneously with somatic cells via a sandwich mechanism

Two models may explain FFWO induced by sperm (Fig. S2C), similarly to what was reported for viruses (Tang et al. 2021). The first model involves an initial fusion of the sperm to a BHK cell expressing JUNO, the transfer of IZUMO1 to the target cell (forming a pseudo-sperm from a pseudo-oocyte) and a subsequent fusion to a new JUNO-expressing BHK (Fig. S2Ci). On the other hand, the second model suggests the fusion of one sperm cell with two JUNO-positive BHK cells, bridging both and forming a continuous syncytium (Fig. S2Cii). We have previously shown that IZUMO1 fades from the sperm after fusion to the fibroblast (Brukman et al. 2023) indicating that this transmembrane protein is transferred to the target cell plasma membrane. Consistent with these findings, IZUMO1 was observed diffusing to the oocyte plasma membrane during in vitro fertilization assays (Satouh et al. 2012). Here we detected IZUMO1 by immunostaining and by a fluorescent reporter and confirmed that it diffuses from the sperm head after its fusion with the BHK cell (Fig. S2A-B), suggesting a function for the fusogenic machinery carried by the sperm during the FFWO process. However, when sperm were first allowed to fuse with one population of green cells before adding the second population of red cells, hybrid syncytia were not observed (Fig. S3A-B). Only when a viral fusogen (VSV-G) was employed or when the two populations of cells were plated before the addition of the sperm, content mixing was induced. These results suggest that it is not possible to uncouple temporarily the sperm-BHK and BHK-BHK fusions, arguing against the model of transfer of fusion proteins and suggesting that sperm are fusing simultaneously to two cells using a sperm sandwich mechanism (Fig. S2Cii).

### Sperm-induced multinucleation as a readout of sperm fertilizing ability

Following the discovery of the ability of sperm to induce cell-cell fusion, we decided to evaluate whether BHK multinucleation could be used as a readout of sperm fusogenic potential. For this purpose, we incubated the sperm in media lacking BSA or Ca^2+^ that do not support capacitation (i.e. a process by which sperm acquires its fusogenic activity; Fig. S4A) (Visconti et al. 1995; Yanagimachi 1988). We found that sperm cells incubated under these conditions failed to fuse to BHK cells, as well as to induce syncytia formation (Fig. 3A-B). Thus, BHK-BHK cell fusion requires fully capacitated sperm. A sperm which is unable to fuse with BHK cells, will not induce multinucleation. Furthermore, to examine whether the extent of cell-cell fusion correlates with the sperm fertilizing ability, we evaluated in parallel the levels of multinucleation and the performance of sperm during *in vitro* fertilization assays. We detected a positive and significant correlation between the syncytia formation and the levels of fertilization, evaluated with complete and denuded oocytes (Fig. 3C-D, Table S1). In contrast, BHK multinucleation did not correlate with the percentage of acrosome reaction of capacitated sperm (Fig. S4B), suggesting that SPICER relies not only on capacitation but on the overall sperm fertilizing potential. Together, these results support the use of this assay as a predictor of sperm fertilizing ability.

### SPICER depends on the activity of IZUMO1

The sperm IZUMO1 protein is essential for sperm-egg interactions via binding to JUNO (Bianchi et al. 2014). More recently, IZUMO1 was shown to induce fusion of cells in a JUNO-independent way using a different domain that is not required for binding to the egg receptor (Brukman et al. 2023). To study whether IZUMO1 is required for the fusion of sperm to BHK cells and for the induction of multinucleation of the fibroblasts, we decided to inhibit IZUMO1 activity using a blocking antibody. For this purpose, we used a monoclonal antibody against IZUMO1 which has been shown to inhibit sperm-egg fusion (Inoue et al. 2013) in our sperm-induced multinucleation assay. We found that the anti-IZUMO1 antibody had the ability to block both sperm-BHK and sperm-induced BHK-BHK fusions even when sperm cells were fully capacitated (Fig. 3A-B). Altogether, our results show that the SPICER assay can determine sperm fertilization potential that depends on the IZUMO-JUNO interactions.

### Cell-cell fusion requires species-specific JUNO

Elegant work on IZUMO1-JUNO interactions among different mammalian species has shown species-specific selectivity between these crucial interactions (Bianchi and Wright 2015). To evaluate the effect of species-specificity of the SPICER assay, we incubated the mouse sperm with BHK cells expressing human JUNO. Under these conditions, we observed lower levels of sperm-BHK fusion and a reduction in BHK multinucleation compared to cells expressing mouse JUNO (Fig. 4A-C). Interestingly, the low activity mediated by human JUNO was still significant compared to the control without JUNO. This confirms the requirement of a species-matching JUNO for the assay; however, it puts in evidence a residual cross-interaction between mouse sperm and human JUNO. In contrast, when hamster JUNO was employed, no differences were observed compared to mouse JUNO (Fig. 4A-C), consistent with the promiscuous nature of the hamster oocytes (Bianchi and Wright 2015; Nakajima et al. 2022; Hanada and Chang 1972).

**Figure 4:**
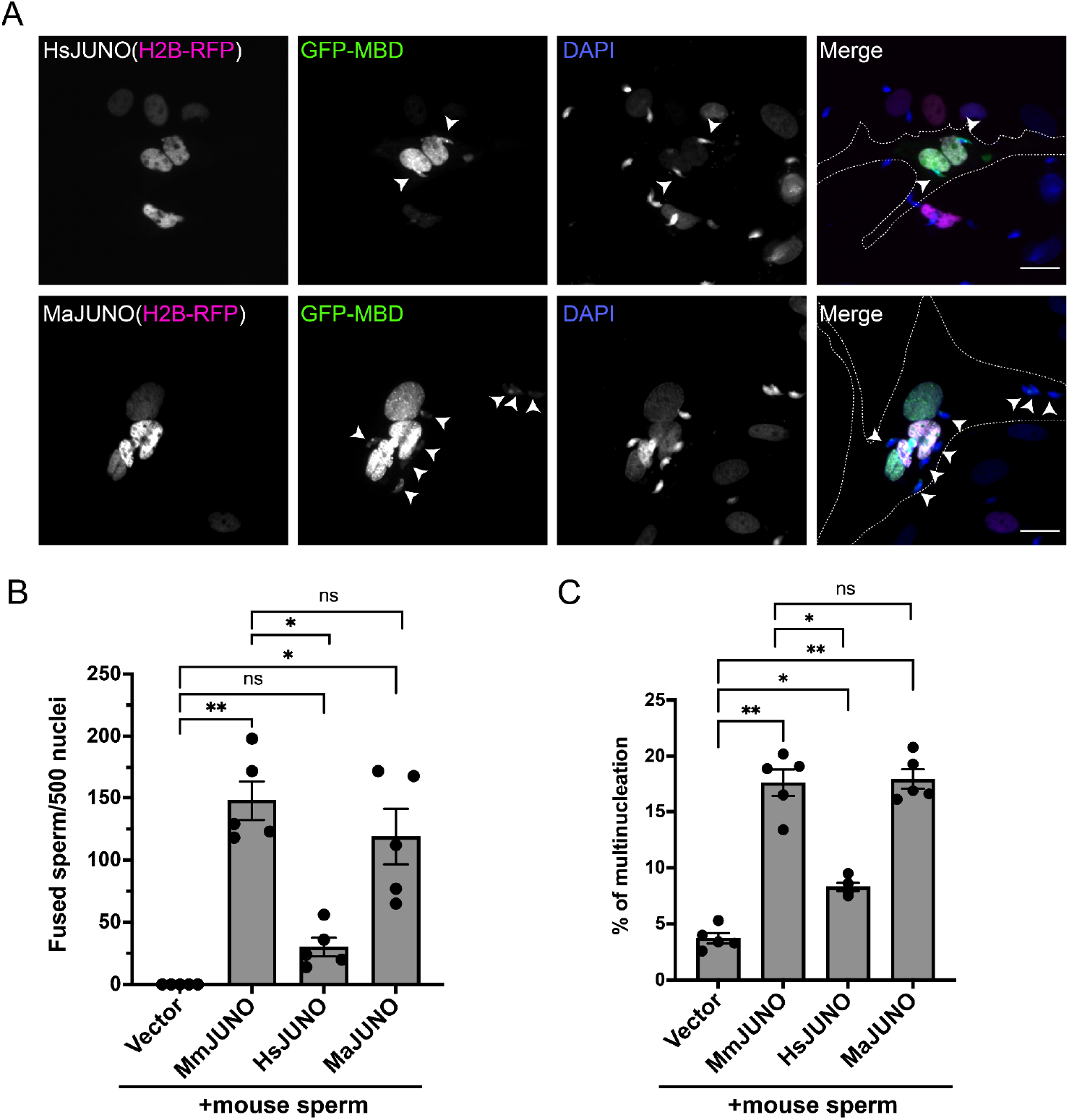
Sperm induce BHK-BHK fusion in a species-specific manner. BHK cells were transfected with pcDNA3.1-EGFP-MBD-nls together pCI::H2B-RFP (empty vector) or pCI::JUNO::H2B-RFP encoding for human JUNO (HsJUNO), mouse JUNO (MmJUNO) or hamster JUNO (MaJUNO), and co-incubated with capacitated mouse sperm. **(A)** Representative images showing H2B-RFP (magenta), GFP-MBD (green), DAPI (blue) channels and the merge. Dotted lines contour cells transfected with human or hamster JUNO with two or eight mouse sperm fused respectively (arrowheads). Scale bar, 20 µm. The number of sperm fused per 500 fluorescent cells **(B)** and the percentage of multinucleation **(C)** were determined. Bar charts showing individual experiment values and means ± SEM of five independent experiments. Comparisons by one-way ANOVA followed by Tukey’s test. *P < 0.05, **P < 0.01, ***P < 0.001.

### Sperm induce the fusion of human epithelial cells

To exclude any BHK cell-specific requirement for SPICER, we tested the epithelial human embryonic kidney 293T (HEK) cells. Due to the smaller size and more rounded shape of these cells that hinder the quantification of multinucleation, we opted to analyze fusion by content mixing. For this, we utilized the Dual Split Protein system (Wang et al. 2014; Ishikawa et al. 2012) which employs two GFP halves expressed in different populations of cells. Upon fusion of these two populations, the reporter self-assembles and the cells become fluorescent (Nakane and Matsuda 2015). The number of green fluorescent cells relative to the number of transfected cells was determined as an indication of fusion levels (Fig. 5A). As observed for the BHK cells, HEK cells bearing JUNO fused between them when they were co-incubated with sperm (Fig. 5B-C). On the contrary, neither HEK cells without JUNO nor HEK expressing JUNO but without sperm showed content mixing (Fig. 5B-C). The fusion kinetics was dependent on the mouse and the co-incubation time reaching a maximum around 4 h (Fig. S5). These results show that SPICER is not restricted to BHK cells or fibroblasts and opens the possibility for adapting this assay to different cell lines and approaches to evaluate cell-cell fusion.

**Figure 5:**
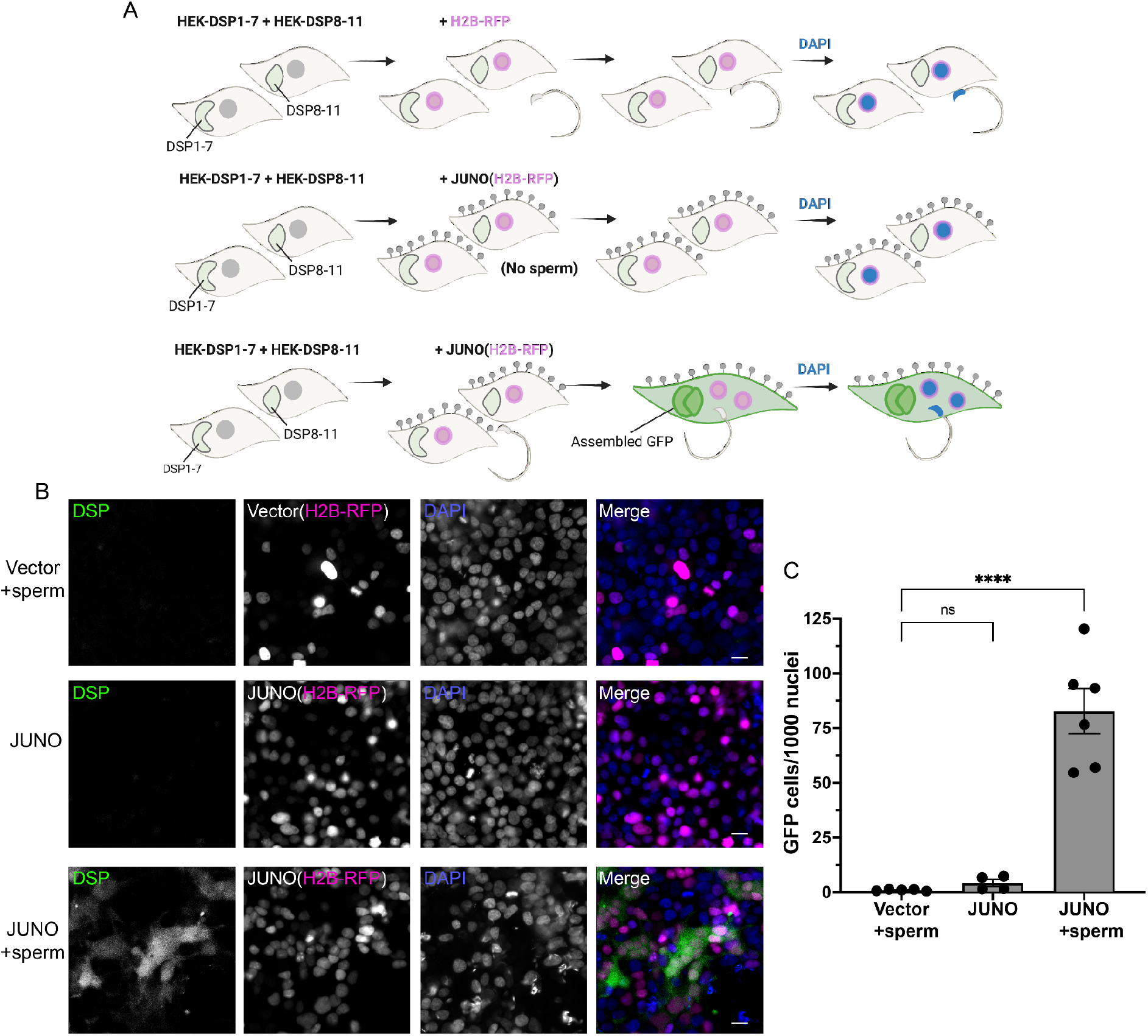
Sperm-induced fusion of human cells evaluated by Dual Split Proteins (DSP) **(A)** Scheme of experimental design. HEK293T cells stably expressing the split GFP (DSP1-7 or DSP8-11) were mixed and transfected with pCI::H2B-RFP or pCI::JUNO::H2B-RFP vectors. When indicated, the cells were later co-incubated with sperm for 4 h, fixed and stained with DAPI. **(B)** Representative images for each treatment. Each separate channel for GFP (assembled GFP in fused cells), H2B-RFP (magenta nuclei) and DAPI staining (blue) are shown. Scale bars, 20 µm. **(C)** Quantification of content mixing experiments. The extent of fusion was determined by counting the number of GFP positive cells per 1,000 nuclei. Bar chart showing individual experiment values (each corresponding to 1,000 to 2,000 nuclei) and means ± SEM of at least four independent experiments. Comparisons by one-way ANOVA followed by Tukey’s test. ns = non-significant, ****P < 0.0001.

## Discussion

In this study, we described a new phenomenon in which sperm cells can induce the fusion of cells ectopically expressing JUNO in culture, resembling the viral-like fusion of cells upon infection. This process is likely mediated by the simultaneous fusion of sperm to adjacent cells and the extent of multinucleation was correlated with the sperm fertilizing potential.

### SPICER occurrence *in vivo*

Apart from being expressed in oocytes, JUNO has been described in lymphoid tissues (Spiegelstein, Eudy, and Finnell 2000), more specifically in regulatory T cells (Yamaguchi et al. 2007). Therefore, we cannot exclude that sperm could bind and fuse to other cell types *in vivo*, and whether it has any physiological or pathological relevance. This fusion cannot occur before acrosomal exocytosis, when IZUMO1 is enclosed into the acrosome (Inoue et al. 2005; Satouh et al. 2012), and therefore, it will be relevant to study sperm fusion to somatic cells within the oviduct of the female, close to the fertilization site (Muro et al. 2016; Hino et al. 2016; La Spina et al. 2016). Furthermore, JUNO was detected in the first polar body (Suzuki et al. 2017) and whether sperm can fuse to it or if sperm fusion to the oocyte can induce the polar body-egg fusion remains unknown.

### Expanding SPICER across species

The assay that we described here can be a potent tool to study cross-fertilization between different species and mechanisms of speciation and evolution, presenting a step beyond biochemical approaches to study IZUMO1-JUNO interactions (Bianchi and Wright 2015). On the other hand, by simply exchanging the JUNO sequence used, this assay could be easily adapted to allow the analysis of the sperm fertilizing ability in different mammalian species, such as humans or cattle, having implications not only for reproductive health but also for biotechnological and agricultural uses. Furthermore, even though IZUMO-JUNO interaction is a specific requirement during sperm-egg fusion of mammals (Grayson 2015), we cannot exclude that alternative configurations, using the right egg receptors (Lu and Ikawa 2022), will be possible in the future to induce somatic cell-cell fusion by sperm of other sexually reproducing organisms. In species where *Izumo1* and *Juno* genes are absent, such as plants and protists, the gamete fusion is promoted by the fusogen HAPLESS 2/GENERATIVE CELL-SPECIFIC 1 (HAP2/GCS1) (Brukman, Li, and Podbilewicz 2022). In other organisms like zebrafish or nematodes, IZUMO1 orthologs were described (Binner et al. 2021; H. Nishimura et al. 2015; Takayama et al. 2021) but, since JUNO is present only in mammals (Vance and Lee 2020; Grayson 2015), it is unclear whether male gametes in other species interact with different receptors on the female gamete. Therefore, it can be speculated, for instance, that the expression of a species-specific Bouncer will be required to activate the fusion of fish sperm (Gert et al. 2023; Herberg et al. 2018) or *Chlamydomonas* FUS1 to mediate the fusion of the algae minus gamete (Pinello, Liu, and Snell 2021; Ferris, Woessner, and Goodenough 1996).

### SPICER as a diagnostic tool

Determining sperm’s fusogenic potential is of great interest for infertility diagnosis of both human and stud animals. Considering that the hamster oocyte penetration (HOP) assay became obsolete, an assay that evaluates the fusion of somatic cells with ectopic expression of JUNO induced by competent sperm (SPICER, Fig. 6) represents a powerful diagnostic tool. SPICER could potentially predict the success chances of ARTs like intrauterine insemination (IUI) or conventional IVF, which require a fusion-competent sperm and if it is necessary to proceed with a more complex technique such as intracytoplasmic sperm injection (ICSI). This is particularly important considering the routine use of the ICSI is under debate as it is associated with a slightly higher risk of adverse outcomes in the progeny (Practice Committees of the American Society for Reproductive Medicine and the Society for Assisted Reproductive Technology 2020).

**Figure 6:**
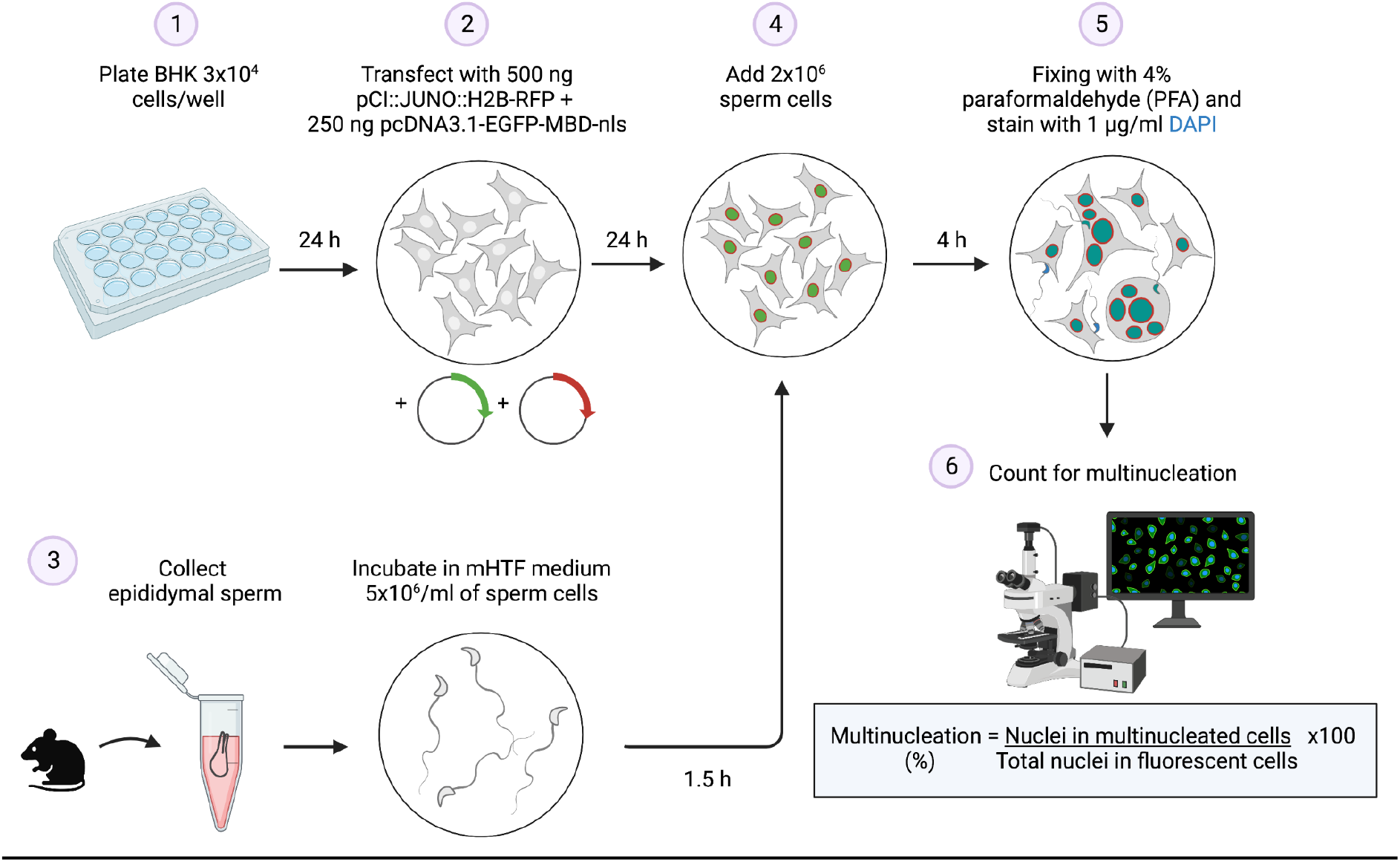
Summary of the SPICER method. Schematic representation of the multinucleation assay employed to determine the sperm fusogenic potential. (1) BHK are seeded on a plate. (2) The cells are transfected with the plasmids encoding for JUNO and EGFP-MBD. (3) Mouse sperm cells are collected and capacitated. (4) The BHK and sperm cells are co-incubated. (5) The cells are washed, fixed and stained with DAPI. (6) Multinucleation levels are quantified as indicated.

Considering our previous work showing that IZUMO1 can mediate cell-cell fusion (Brukman et al. 2023) and that significantly lower levels of this protein were detected in the sperm cells from patients with total fertilization failure (Enoiu et al. 2022), we expect that the SPICER assay will be able to resolve some unexplained cases of male infertility that result from loss of function of components of the fusion machinery. In this context, other proteins that are essential for gamete fusion, such as SPACA6, TMEM95, TMEM81, FIMP, SOF1 and DCST1/2 (Barbaux et al. 2020; Noda et al. 2020; Lorenzetti et al. 2014; Lamas-Toranzo et al. 2020; Fujihara et al. 2020; Inoue, Hagihara, and Wada 2021; Noda et al. 2022; Elofsson et al. 2023; Deneke et al. 2023), might be transferred to the somatic cells together with IZUMO1 during sperm-induced cell-cell fusion. Finally, considering that antibodies against IZUMO1 were detected on the sera of infertile women (Clark and Naz 2013) and here we observed that a monoclonal antibody against this protein was able to block sperm-induced syncytia, SPICER could serve as tool to readily analyze cases of immuno-infertility. For all these cases, in-depth studies utilizing human sperm from healthy donors and infertile patients will be necessary.

### Further applications for SPICER

In addition to the potential uses of SPICER for diagnosis of infertility, this assay could be used to evaluate potential sperm donors and animal studs. Moreover, our SPICER tool has the potential to aid screening for compounds that enhance fertilization (new fertility treatments) or that block gamete interactions (Stepanenko et al. 2022), or to easily determine the effect of genetic variations of JUNO (Allingham and Floriano 2021; Takaiso et al. 2016). SPICER could also facilitate studies aimed at resolving standing enigmas of sperm-egg fusion (Deneke and Pauli 2021; Yanagimachi 2022). This includes the role of IZUMO1 and other potential fusogens as well as additional cellular pathways that are known to influence fusion in different systems, including the action of the cytoskeleton and molecular motors (Shilagardi et al. 2013; Yang et al. 2017; Zhang et al. 2017), specific lipids (e.g. phosphatidylserine) (Rival et al. 2019; van den Eijnde et al. 2001; Verma et al. 2018; Abay et al. 2017), calcium signaling (Tsuchiya et al. 2018; Earles et al. 2001), phosphorylation cascades (Eigler et al. 2021) and more (Brukman et al. 2019). Finally, whether other cell types rather than sperm, such as muscle cells or osteoclasts, can induce syncytia formation after ectopic fusion remains an open question, with important biological implications.

### Limitations of the study

The main caveat of our assay is that it cannot discriminate between defects in sperm docking or gamete fusion, as a poor performance in SPICER could be explained by the altered interaction of IZUMO1 and JUNO or by a reduced capacity of IZUMO1 (and IZUMO1 partners in the sperm) to induce membrane merger. Moreover, simultaneous measurements of the levels of spontaneous acrosome reaction will be needed to exclude a faulty capacitation. Alternatively, capacitation could be further stimulated pharmacologically using a Ca^2+^ ionophore (Tateno et al. 2013). In this context, further analysis may be required to refine the characterization of defective sperm. As noted before, future validations using human sperm will be necessary for the establishment of an assay that could be used in the clinics.

## MATERIALS AND METHODS

### RESOURCES TABLE

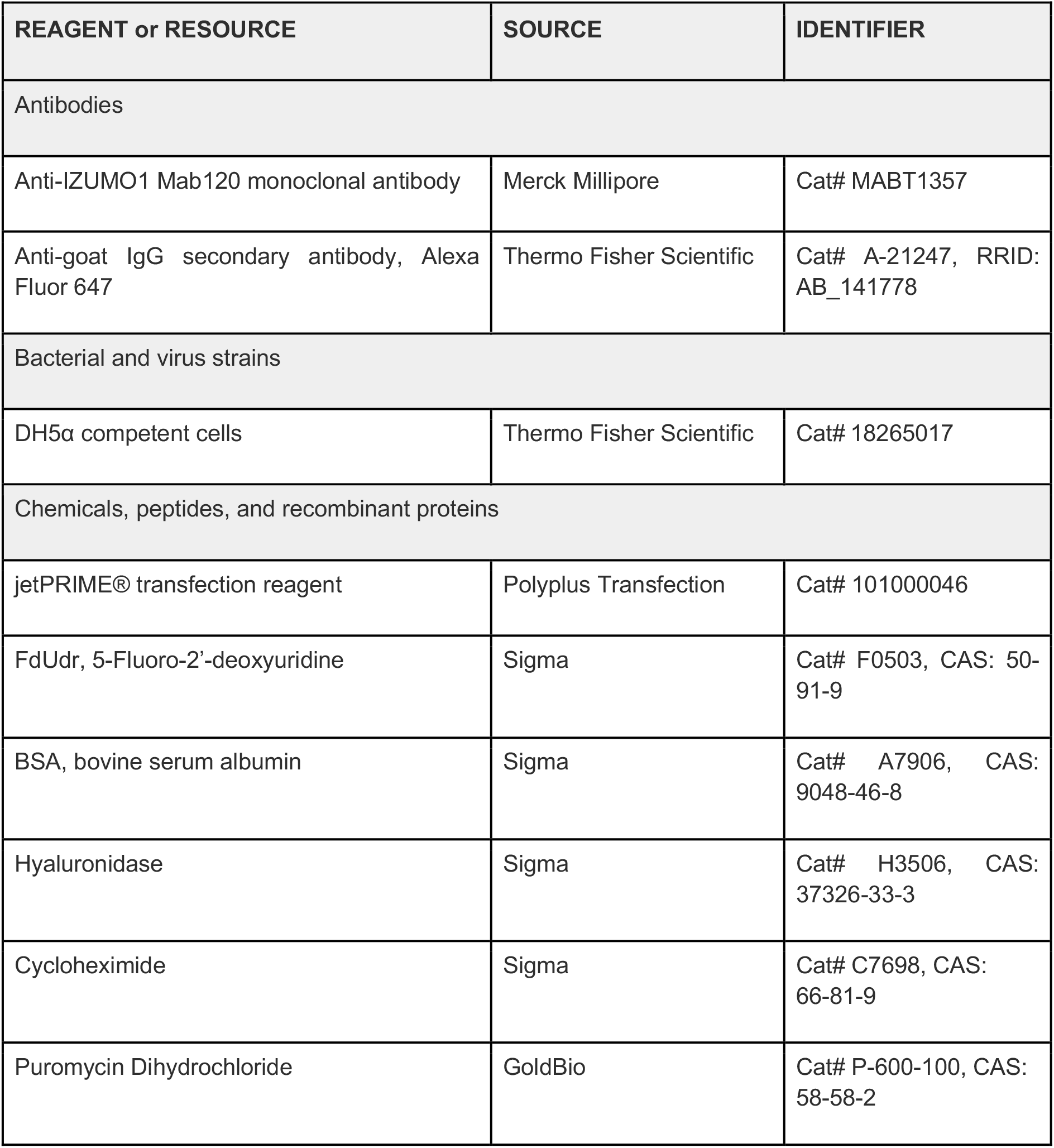
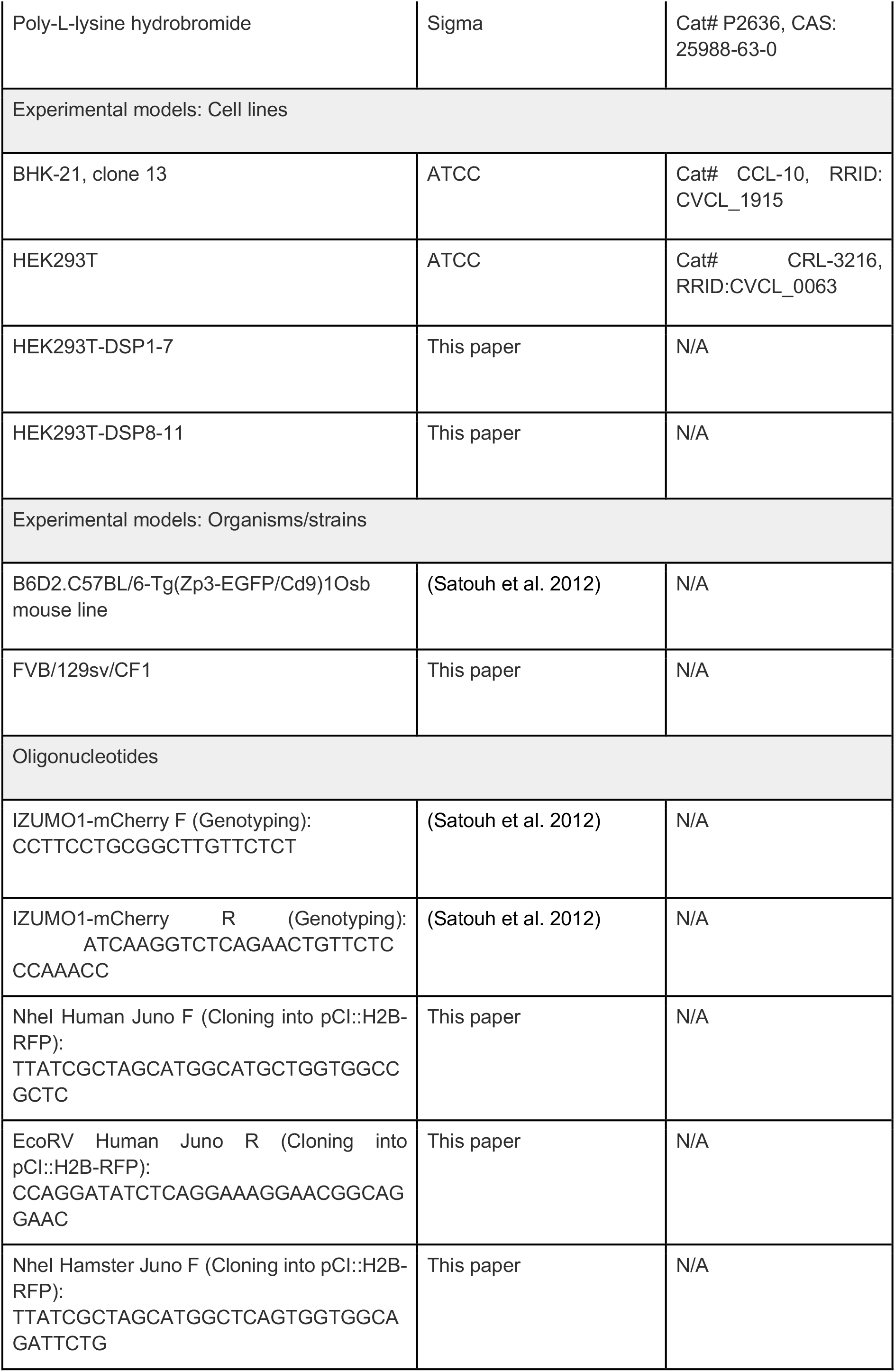
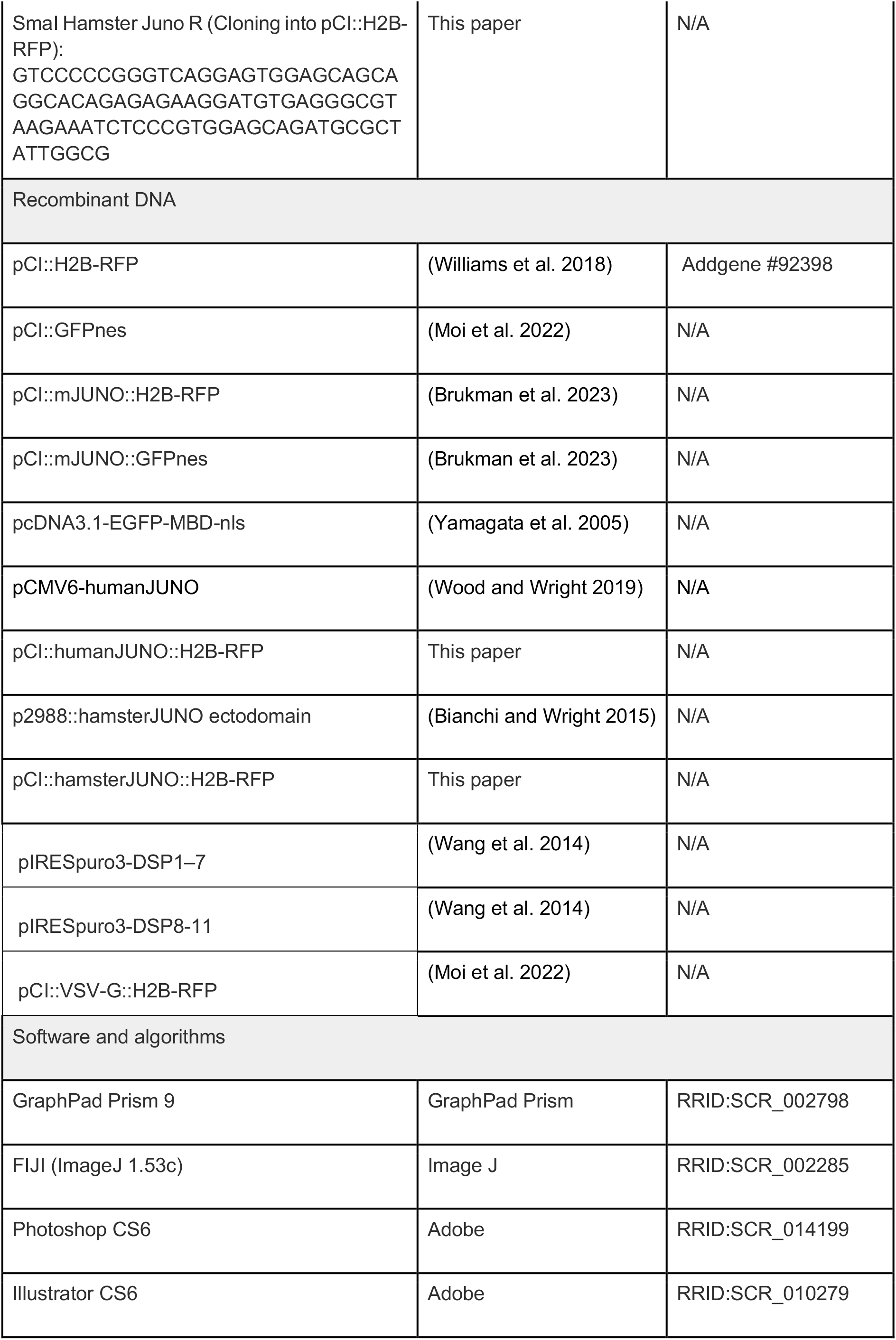
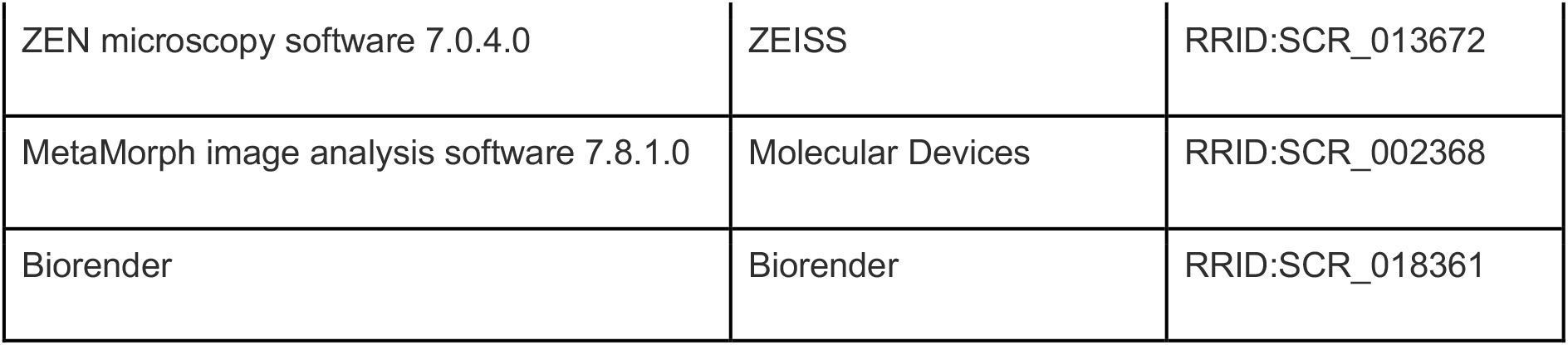

### RESOURCE AVAILABILITY

#### Lead contact

Further information and requests for resources and reagents should be directed to and will be fulfilled by the lead contact, Benjamin Podbilewicz.

#### Materials availability

Plasmids or cell lines generated in this study are available upon request. Signing a materials transfer agreement (MTA) may be required.

#### Data and code availability

Any additional information required to reanalyze the data reported in this paper is available from the lead contact upon request.

### EXPERIMENTAL MODEL AND SUBJECT DETAILS

#### Animals

All animal studies were approved by the Committee on the Ethics of Animal Experiments of the Technion - Israel Institute of Technology. In this study, we used wild-type male mice from a FVB/129sv/CF1 mixed background and B6D2-Tg(Izumo1-mCherry) mouse line (Satouh et al. 2012). Animals were bred and housed in the Technion animal facility under specific pathogen–free conditions with *ad libitum* access to food and water. The primers used for genotyping Izumo1-mCherry are outlined in the Key Resources Table. In all cases, male mice between 3 and 6 months old were used for the experiments.

#### Cell lines and DNA transfection

BHK cells (Cat# CCL-10; ATCC, RRID: CVCL_1915) and HEK293T cells (Cat# CRL-3216; ATCC, RRID: CVCL_0063) were grown and maintained in DMEM containing 10% FBS. Cells were cultured at 37°C in 5% CO_2_. Plasmids were transfected into cells using 2 μl jetPRIME (PolyPlus-transfection) per µg of DNA in 100 μl of reaction buffer for every ml of medium. HEK293T cells stable lines for Dual Split Proteins (DSP) 1-7 and 8-11 were prepared by transfecting pIRESpuro3-DSP1–7 and pIRESpuro3-DSP8-11 respectively and selecting with 2 µg/ml of puromycin for 10–13 days as previously described (Wang et al. 2014).

### METHOD DETAILS

#### Sperm collection and capacitation

Sperm were recovered by incising the cauda epididymis, obtained from adult male mice, in 300 μl of mHTF medium (Kito et al. 2004) supplemented with 4 mg/ml of BSA. The sperm were diluted in fresh medium to a concentration of 5 × 10^6^ cells/ml and incubated for 90 min at 37°C and 5% CO_2_ to induce capacitation.

#### Sperm-to-BHK fusion and multinucleation

BHK cells were grown on 24-well glass bottom tissue-culture plates. 24 h after plating, cells were transfected with 0.25 µg pcDNA3.1-EGFP-MBD-nls plasmids and 0.5 µg of either pCI::H2B-RFP or pCI::JUNO::H2B-RFP. 24 h after transfection, 2 × 10^6^ capacitated wild-type sperm cells in mHTF were added to each well and co-incubated with the BHK cells for 4 h at 37°C and 5% CO_2_. After one wash with PBS, the cells were fixed with 4% PFA in PBS and stained with 1 µg/ml DAPI. Micrographs were obtained using wide-field illumination using an ELYRA system S.1 microscope (Plan-Apochromat 20× NA 0.8; Zeiss). Multinucleation percentage was determined as the ratio between the number of nuclei in multinucleated cells (NuM) and the total number of nuclei in fluorescent cells (NuF), as follows: % of multinucleation = (NuM/NuF) × 100. 500 nuclei (NuF) were counted in each independent repetition (experimental point). In some cases, the number of sperm fused was determined by evaluating the transfer of EGFP-MBD-nls signal from the BHK cell to the sperm nuclei (number of fused sperm/500 BHK cells; independently of the number of nuclei within each cell). In some experiments, sperm cells obtained from transgenic mice expressing IZUMO1-mCherry were employed to analyze IZUMO1 localization.

#### *In vitro* fertilization

Ovulated oocytes were obtained from females previously treated with an i.p. injection of pregnant mare serum gonadotropin (5 IU; #HOR-272, Prospec), followed by an i.p. injection of human chorionic gonadotropin (5 IU, #CG5; Sigma-Aldrich) 48 h later. Cumulus–oocyte complexes (COCs) were collected from the ampullae of induced females 12–15 h after human chorionic gonadotropin administration in mHTF medium. COCs were inseminated with 5 × 10^3^ capacitated sperm and co-incubated in capacitation media for 3 h at 37°C and 5% CO_2_. Then, the oocytes were washed, stained with 10 µg/ml Hoechst 33342 (Sigma) and observed using wide-field illumination using an ELYRA system S.1 microscope (Plan-Apochromat 20× NA 0.8; Zeiss). Eggs were considered fertilized when at least one decondensing sperm nucleus or two pronuclei were observed in the egg cytoplasm. For fusion quantification, oocytes were denuded from the cumulus and the ZP by sequential treatment with 0.3 mg/ml hyaluronidase (H3506; Sigma-Aldrich) and acid Tyrode solution (pH 2.5; (Nicolson, Yanagimachi, and Yanagimachi 1975)). ZP-free eggs were inseminated with 10^3^ capacitated sperm and co-incubated for 1 h. Then, the eggs were processed as above and the number of decondensing sperm nuclei per oocyte was scored.

#### Content mixing assay using different colors

BHK cells at 70% confluence in 35-mm plates were transfected with 1 µg pCI::H2B-RFP, pCI::GFPnes, pCI::JUNO::H2B-RFP or pCI::JUNO::GFPnes. 4 h after transfection, the cells were washed four times with DMEM with 10% serum, four times with PBS, and detached using Trypsin (Biological Industries). The cells were collected, resuspended in DMEM with 10% serum, and counted. Equal amount of H2B-RFP and GFPnes cells (1.25 × 10^5^ each) were mixed and seeded on glass-bottom plates (12-well black, glass-bottom #1.5H; Cellvis) and incubated at 37°C and 5% CO_2_. 18 h after mixing, 4 × 10^6^ capacitated wild-type sperm cells in mHTF were added to the BHK cells and co-incubated for 4 h after which they were washed with PBS, fixed with 4% PFA in PBS and stained with 1 µg/ml DAPI. Micrographs were obtained using wide-field illumination using an ELYRA system S.1 microscope (Plan-Apochromat 20× NA 0.8; Zeiss). The percentage of mixing was defined as the ratio between the nuclei in mixed cells (NuM) and the total number of nuclei in mixed cells and fluorescent cells whose cell bodies were in contact and did not fuse (NuC), as follows: % of mixing = (NuM/[NuM + NuC]) × 100. 1,000 nuclei (NuM + NuC) were counted in each independent repetition (experimental point).

#### Content mixing assay in two steps

BHK cells grown on 12-well glass bottom tissue-culture plates were transfected with 1 µg of either pCI::GFPnes or pCI::JUNO::GFPnes. 24 h later, the cells were thoroughly washed with DMEM with 10% serum, and, when indicated, 4 × 10^6^ capacitated wild-type sperm cells in mHTF were added to the BHK cells, and co-incubated for 1 h. In some cases, the sperm were removed by washing three times with DMEM. In parallel, BHK grown in 35-mm plates that were transfected with 1 µg pCI::JUNO::H2B-RFP or pCI::VSV-G::H2B-RFP the day before were washed and detached using 0.05% EDTA solution. These cells were added in a 1:1 ratio to the BHK cells previously incubated or not with the sperm, as indicated. 18 h later, the cells were fixed with 4% PFA in PBS, stained with 1 µg/ml DAPI, and content mixing was evaluated as explained above. For activating VSV-G activity, 1 h before fixing a 5-minute incubation at pH 5.5 buffer was performed (Moi et al. 2022).

#### Content mixing assay using the Dual Split Protein

Equal numbers of HEK293T cells stably expressing DSP1-7 or DSP8-11 were mixed (1.25 × 10^5^ each) and seeded on glass-bottom plates (12-well black, glass-bottom #1.5H; Cellvis) pre-treated with 20 μg/ml of Poly-L-lysine. 24 h later, the cells were transfected with 1 µg pCI::H2B-RFP or pCI::JUNO::H2B-RFP. 18 h after transfection, 4 × 10^6^ capacitated wild-type sperm cells in mHTF were added to the HEK293T cells and co-incubated for 4 h after which they were washed with PBS, fixed with 4% PFA in PBS and stained with 1 µg/ml DAPI. In addition, a time course experiment was conducted where the cells were fixed at different time points. Micrographs were obtained as above, using wide-field illumination using an ELYRA system S.1 microscope (Plan-Apochromat 20× NA 0.8; Zeiss). The number of GFP positive cells per 1,000 nuclei was determined. Between 1,000 to 2,000 red nuclei were counted in each independent repetition (experimental point).

#### Inhibition of sperm-induced cell-cell fusion

Different conditions were tested to evaluate their effect on sperm-induced cell-cell fusion. Non-capacitated sperm were incubated in medium mHTF lacking Ca^2+^ or BSA for 90 min at 37°C and 5% CO_2_. In other cases, capacitated sperm were mixed before their addition to the BHK cells with 1 µg/µl of anti-IZUMO1 antibody (Inoue et al. 2013), 20 µM of the cell cycle inhibitor FdUdr (Valansi et al. 2017) or 200 µg/ml of the inhibitor of protein synthesis cycloheximide (Wengler 1975).

#### Evaluation of acrosome reaction

The extent of acrosome reaction was evaluated by Coomassie brilliant blue staining as previously described (Busso et al. 2007). Briefly, the sperm cells were fixed in 4 % paraformaldehyde in PBS for 15 min at room temperature, washed with 0.1 M ammonium acetate (pH 9) by centrifugation, mounted on slides, and air dried. Slides were successively immersed 5 min in water, 5 min in ice-cold methanol, 5 min in water, and 2 min in 0.22% Coomassie brilliant blue solution (50% methanol and 10% acetic acid). After washing with water, the samples were mounted and observed under a light microscope (X200). Sperm were scored as acrosome-intact when a bright blue labeling was observed in the dorsal region of the head or as acrosome-reacted when no staining was observed. For each condition, 1,000 sperm were counted.

#### Time-lapse Imaging of sperm-induced cell-cell fusion

BHK cells were grown on 35 mm glass bottom tissue-culture plates (Greiner Bio-one) and, 24 h after plating, cells were transfected with 1.5 µg pCI::JUNO::GFPnes. 24 h after transfection, time-lapse images of the cells were imaged before and after adding 5 × 10^6^ capacitated wild-type sperm cells in mHTF. Images of the cells were acquired every 3 min for 1 h to record cell-to-cell fusion, using a spinning disc confocal microscope (CSU-X; Yokogawa Electric Corporation) with an Eclipse Ti inverted microscope and a Plan-Apochromat ×20 (NA, 0.75; Nikon) objective. Images were obtained using an iXon3 EMCCD camera (ANDOR) through MetaMorph (Molecular Devices, version 7.8.1.0). Images in differential interference contrast and green channels were recorded.

#### Immunofluorescence

The localization of IZUMO1 was determined by immunostaining. Briefly, after fixation cell were permeabilized with 0.1% Triton X-100 in PBS and incubated with anti-IZUMO1, clone Mab120 (1:500, Cat# MABT1357; Merck Millipore) followed by the secondary antibody Alexa Fluor 647 goat anti-rat (1:500, Cat# A-21247; Thermo Fisher Scientific, RRID: AB_141778). Later, the nuclei were stained with 1 µg/ml DAPI and micrographs were obtained using wide-field illumination using an ELYRA system S.1 microscope (Plan-Apochromat 20× NA 0.8; Zeiss) with an EMCCD iXon camera (Andor) through ZEN microscopy software 7.0.4.0 (RRID: SCR_013672; Zeiss).

### QUANTIFICATION AND STATISTICAL ANALYSIS

#### Statistics and data analysis

Results are shown as means ± SEM. For each experiment, at least three independent biological repetitions were performed. The significance of differences between the averages were analyzed using one-way ANOVA and Pearson’s analysis was used to assess the correlations, as described in the legends (GraphPad Prism 9, RRID: SCR_002798). Figures were prepared with Photoshop CS6 (Adobe, RRID: SCR_014199), Illustrator CS6 (Adobe, RRID: SCR_010279), BioRender.com (RRID: SCR_018361), and FIJI (ImageJ 1.53c, RRID: SCR_002285).

## Acknowledgements

We thank Gavin Wright and Enrica Bianchi (University of York, York, UK) for the plasmids encoding for human and hamster JUNO, Kazuo Yamagata (Kindai University, Higashiosaka City, Osaka, Japan) for the EGFP-MBD-NLS plasmid, Masahito Ikawa (Osaka University, Osaka, Japan) for the IZUMO1-mCherry transgenic mouse line, and Zene Matsuda (University of Tokyo, Japan) for the DSP plasmids. Finally, we thank Dan Cassel (Technion-Israel Institute of Technology), Leonid Chernomordik (NICHD/DIR, NIH, US) and the members of Podbilewicz Lab for critically reading the manuscript.

This project has received funding from the Israel Science Foundation (257/17, 2462/18, 2327/19, and 178/20 to B. Podbilewicz), the European Union’s Horizon 2020 research and innovation program under the Marie Skłodowska-Curie Actions (844807 to N.G. Brukman) and Direccion General de Asuntos del Personal Academico, Programa de Estancias de Investigacion (PREI), UNAM, Mexico City (B. Podbilewicz).

## Author contributions

B.P. conceived the study. N.G.B. and C.V. designed and conducted experiments, and processed and analyzed the data supervised by B.P.; N.G.B. wrote the initial draft of the manuscript. All authors participated in discussions of results and manuscript editing.

## Supplementary Information

**Figure S1:**
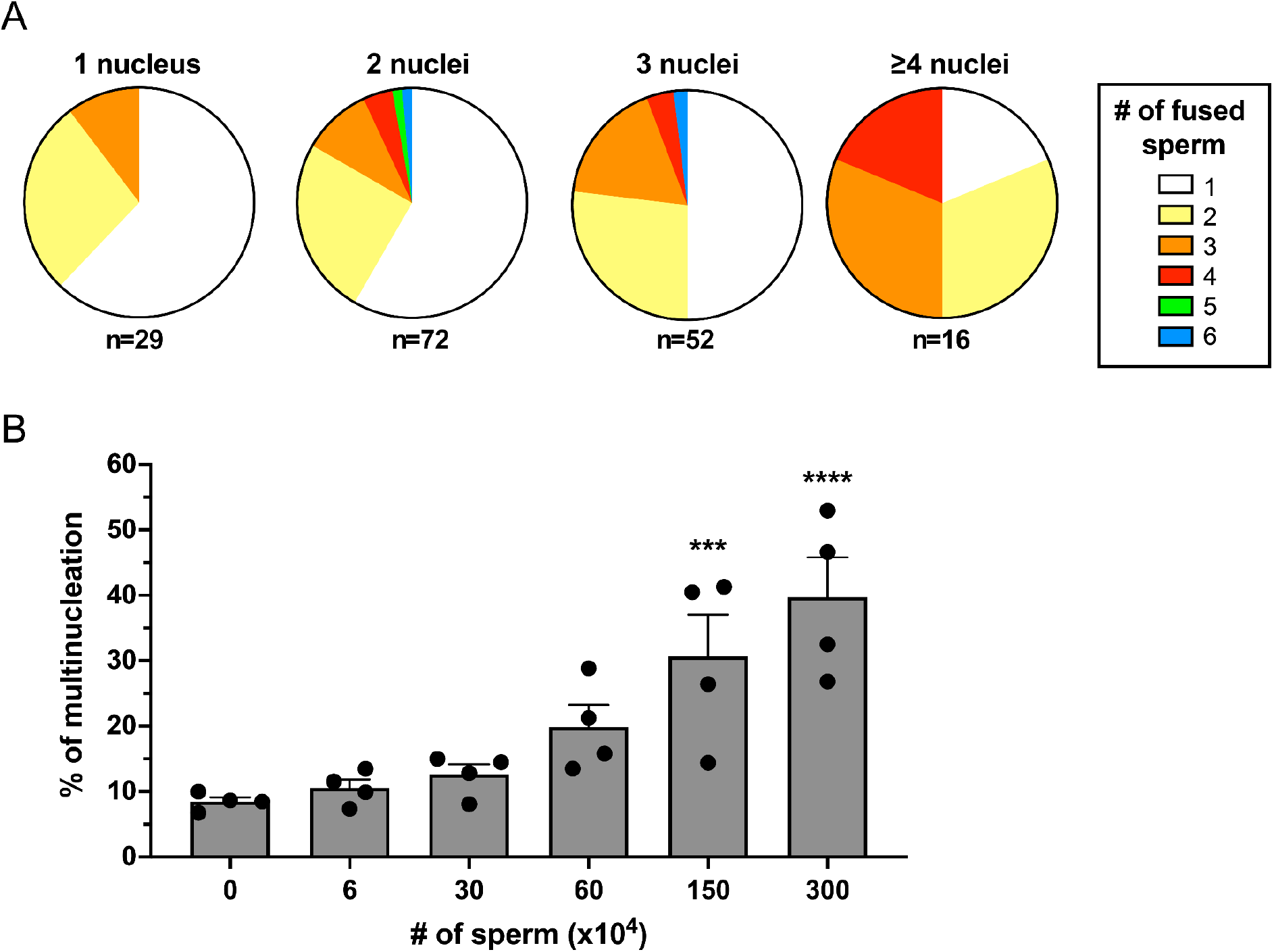
Effect of sperm number on syncytia formation. BHK cells were transfected with pCI::JUNO::H2B-RFP vectors and pcDNA3.1-EGFP-MBD-nls co-incubated with capacitated sperm for 4 h. **(A)** The distribution of BHK cells with 1, 2, 3 or 4 nuclei presenting different amounts of sperm fused to them. **(B)** Different amounts of sperm were added to the cells. The percentage of multinucleation was defined as the ratio between the nuclei in multinucleated cells (NuM) and the total number of nuclei in fluorescent cells (NuF), as follows: % of multinucleation = (NuM/NuF) × 100. We show individual data and means ± SEM of four independent experiments. The number of nuclei counted per experiment and per treatment was 1,000. Comparisons were made with one-way ANOVA followed by Dunnett’s test against the control without sperm (0). ***P < 0.001, ****P < 0.0001.

**Figure S2:**
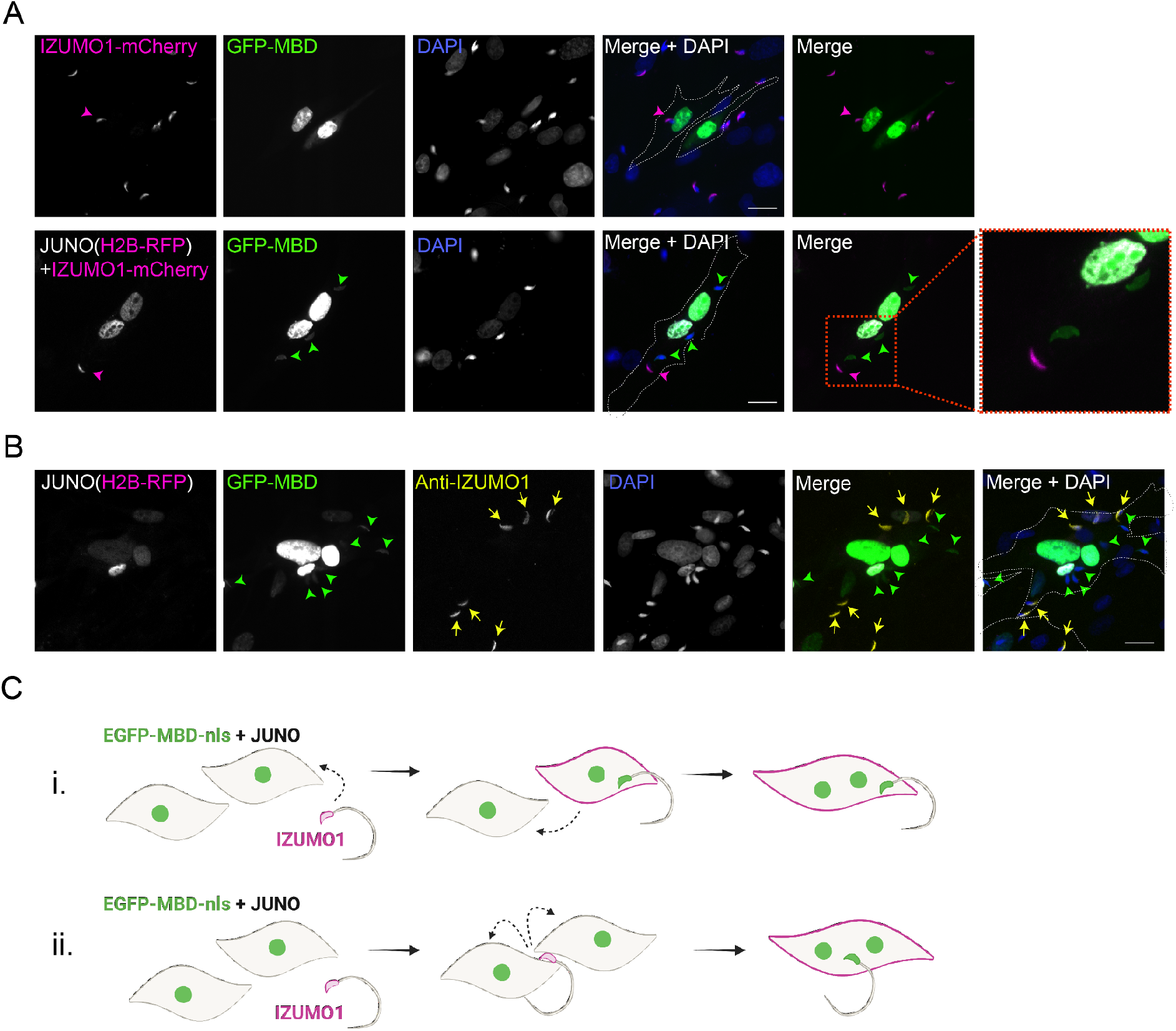
IZUMO1 diffuses into the target cell upon sperm fusion. **(A)** BHK were transfected with pcDNA3.1-EGFP-MBD-nls alone or together with pCI::JUNO::H2B-RFP and co-incubated with sperm obtained from transgenic mice expressing IZUMO1-mcherry. Representative images showing H2B-RFP and Izumo1-mCherry (magenta), GFP-MBD (green), DAPI (blue) channels and the merge with and without DAPI. Dotted lines contour relevant cells. Scale bar, 20 µm. Magenta arrowheads point to unfused sperm (GFP negative) presenting IZUMO1-mCherry signal while green arrowheads point to fused sperm (GFP positive) without Izumo1-mCherry signal. **(B)** BHK were transfected with pcDNA3.1-EGFP-MBD-nls together with pCI::JUNO::H2B-RFP and co-incubated with wild-type mouse sperm. After fixation, an immunostaining against IZUMO1 was performed. Representative images showing H2B-RFP (magenta), GFP-MBD (green), IZUMO1 staining (yellow), DAPI (blue) channels and the merge with and without DAPI. Dotted lines contour relevant cells. Scale bar, 20 µm. Yellow arrows point to unfused sperm (GFP negative) presenting IZUMO1 signal while green arrowheads point to fused sperm (GFP positive) without IZUMO1 signal. **(C)** Models for sperm-induced cell-cell fusion. In general, this phenomenon resembles the so-called “fusion from without” in which viruses can induce the fusion of the target cells (Falke, Knoblich, and Müller 1985; Tang et al. 2021; Edwards and Brown 1986). i-One sperm fuses with a BHK, IZUMO1 is transferred by diffusion and mediates the fusion to another cell. ii-One sperm fuses to two BHK bridging them and inducing the formation of a syncytium.

**Figure S3:**
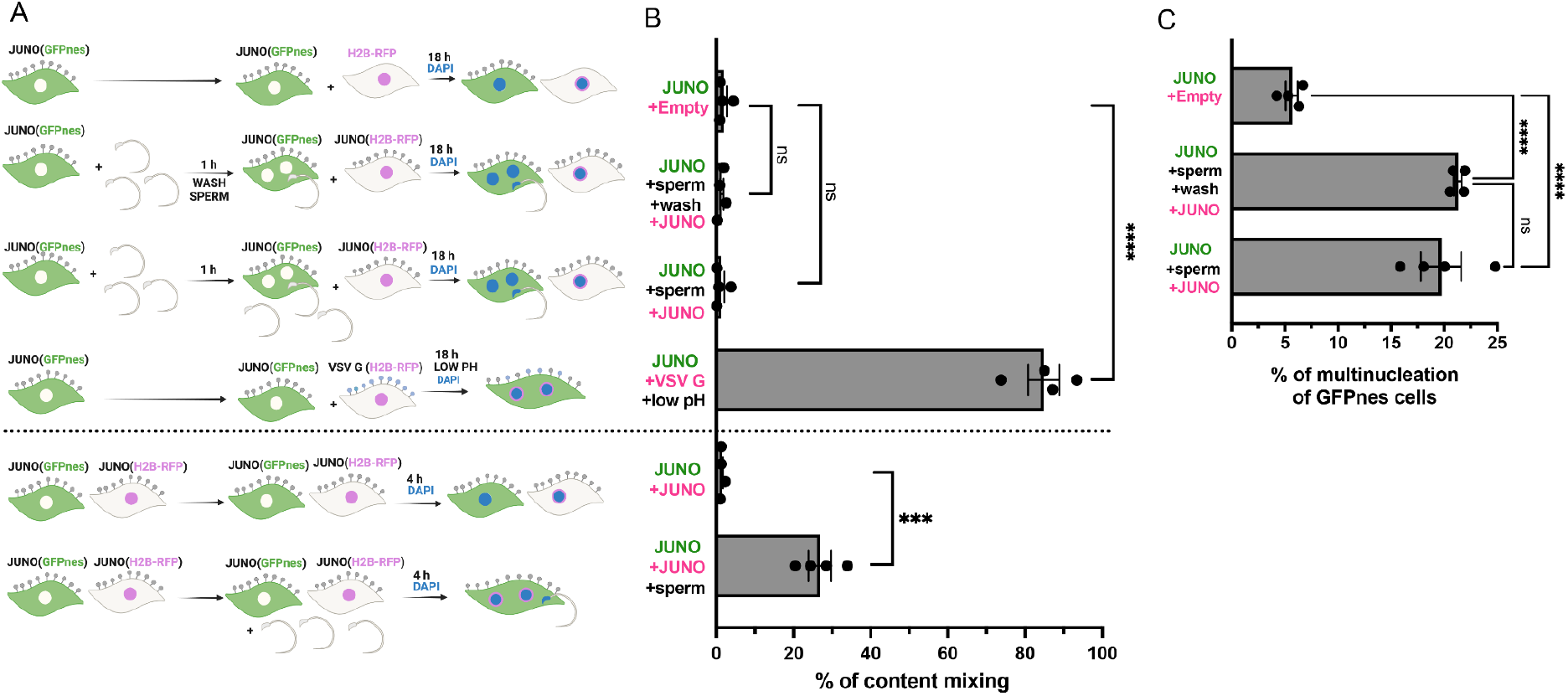
The timing of sperm fusion to somatic cells is critical for content mixing. **(A)** Scheme of experimental design. BHK cells transfected with pCI::JUNO::GFPnes were incubated or not with sperm for 1 h. When indicated the sperm were washed and cells previously transfected with pCI::H2B-RFP, pCI::JUNO::H2B-RFP, or pCI::VSV-G::H2B-RFP were added. After 18 h, the cells were fixed and stained with DAPI. To activate VSV-G a treatment with low pH was done. In parallel, the regular mixing assay was performed in which BHK cells transfected with pCI::JUNO::GFPnes or pCI::JUNO::H2B-RFP were previously mixed and then co-incubated (or not) with sperm for 4 h, fixed and stained with DAPI. **(B)** Quantification of content mixing experiments. The percentage of mixing was defined as the ratio between the nuclei in mixed cells (NuM) and the total number of nuclei in mixed cells and fluorescent cells whose cell bodies are in contact that did not fuse (NuC), as follows: % of mixing = (NuM/[NuM + NuC]) × 100. Bar chart showing individual experiment values (each corresponding to 1,000 nuclei) and means ± SEM of four independent experiments. For the 18 h mixing: comparisons by one-way ANOVA followed by Dunnett’s test against the negative control. For the 4 h mixing: comparison by t-Student’s test. ns = non-significant, ***P < 0.001, ****P < 0.0001. **(C)** Multinucleation was quantified for the pCI::JUNO::GFPnes-transfected cells for the indicated conditions. We show individual data and means ± SEM of four independent experiments. Comparisons by one-way ANOVA followed by Tukey’s test. ns = non-significant, ****P < 0.0001.

**Figure S4:**
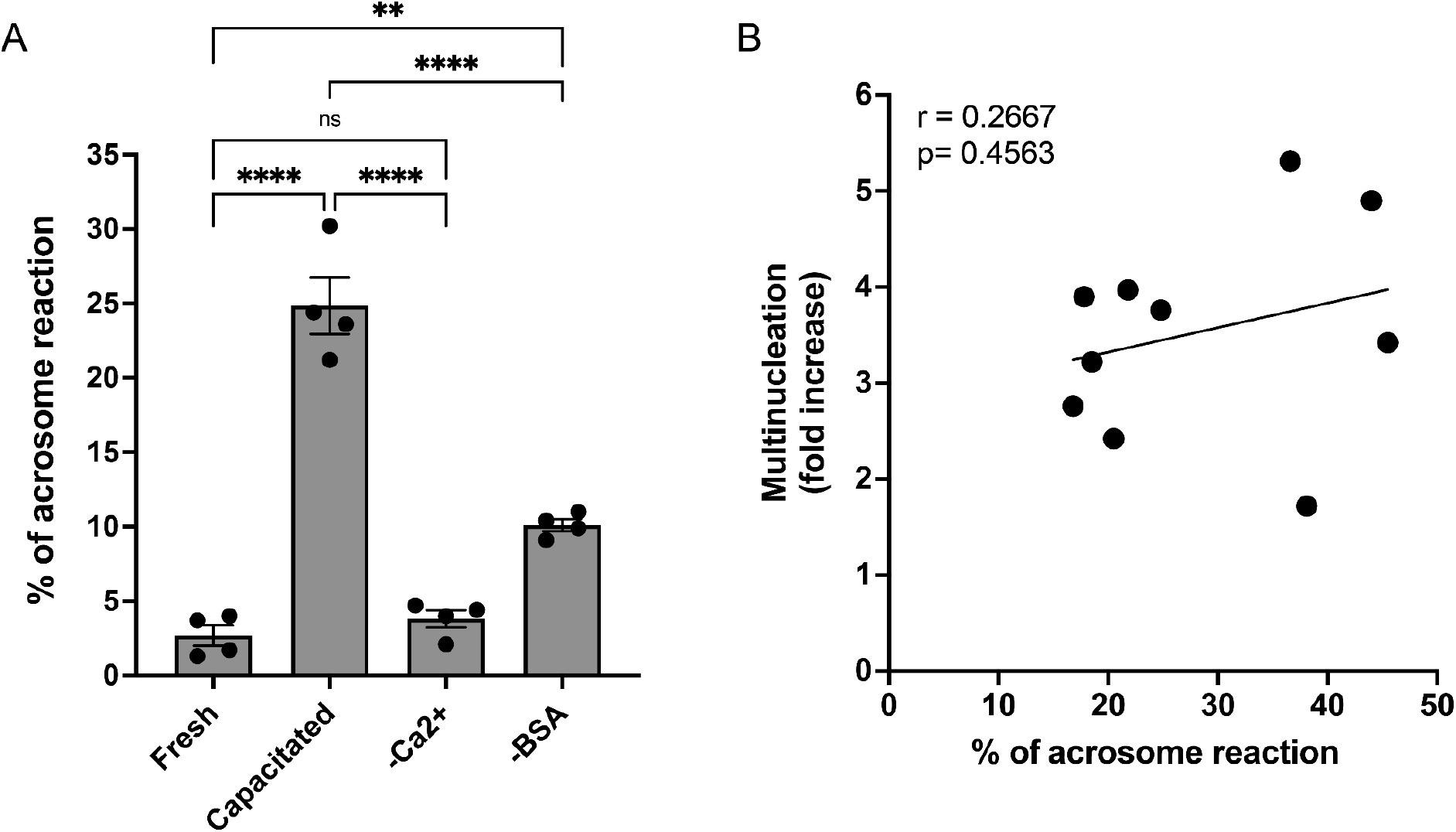
Acrosome reaction is reduced in non-capacitated sperm, but does not correlate with SPICER levels in capacitated conditions. **(A)** Percentage of acrosome reaction of fresh sperm, capacitated sperm, and sperm incubated in media lacking calcium (-Ca^2+^) or bovine serum albumin (-BSA). Comparisons by one-way ANOVA followed by Tukey’s test. ns = non-significant, **P < 0.01, ****P < 0.0001. **(B)** Multinucleation levels relative to the control without sperm as a function of the percentage of acrosome reaction after capacitation. Each dot corresponds to a different mouse. The Pearson’s coefficient ‘r’ and the significance are included.

**Figure S5:**
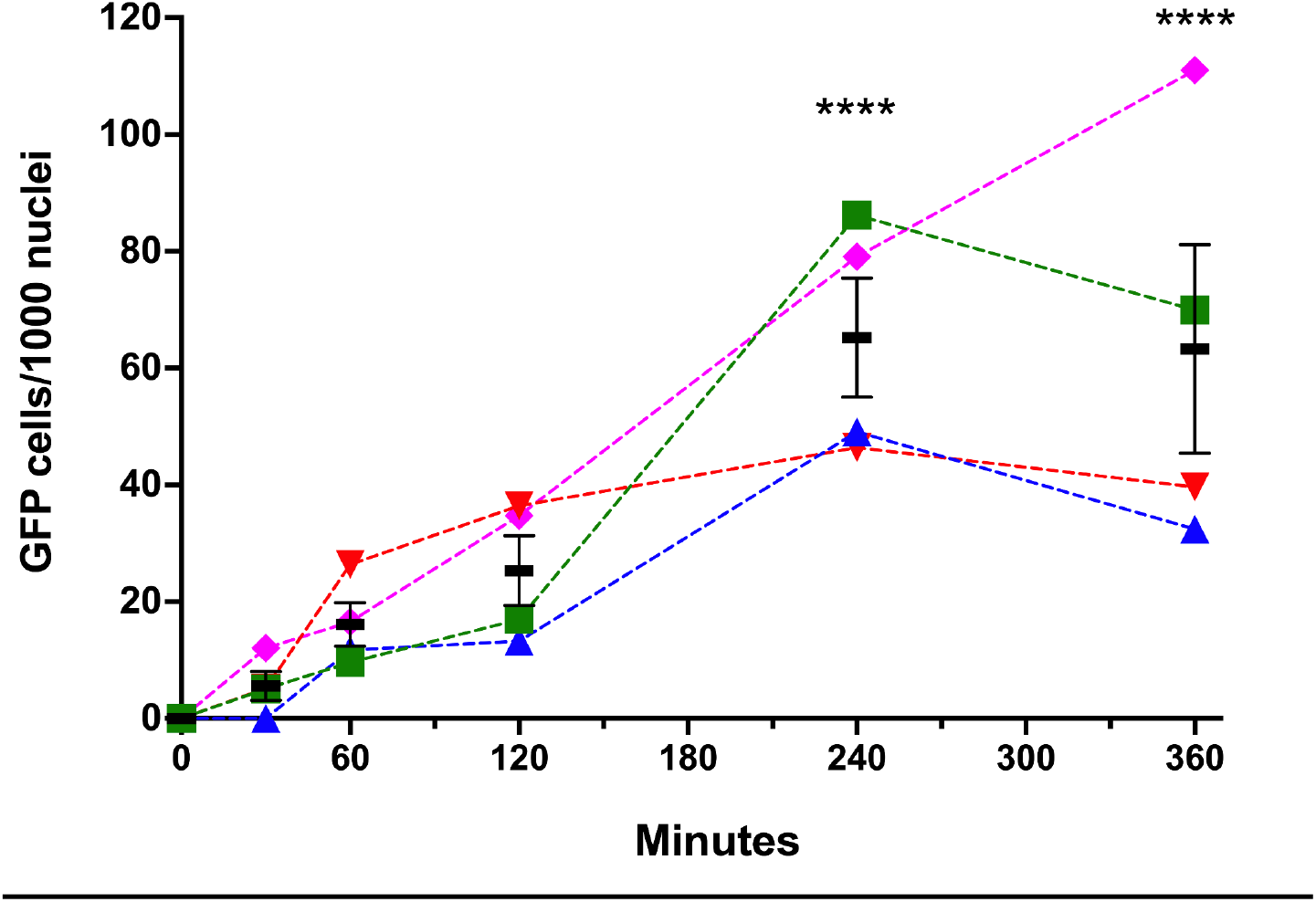
Time course of sperm-induced fusion of human cells evaluated by Dual Split Proteins (DSP) HEK293T cells stably expressing the split GFP (DSP1-7 or DSP8-11) were mixed and transfected with pCI::JUNO::H2B-RFP vectors. The cells were later co-incubated with capacitated sperm for 0, 30, 60, 120, 240, and 360 min, fixed, and stained with DAPI. The extent of fusion was determined by counting the number of GFP-positive cells per 1,000 nuclei. Results from four different mice are plotted in different colors and symbols. The means ± SEM of the four independent experiments are shown in black. Comparisons by one-way ANOVA followed by Dunnett’s test against 0 min. ****P < 0.0001.

**<u>Movie S1:</u>** Time-lapse experiment using spinning disk confocal microscopy showing the fusion of two cells expressing JUNO and GFPnes after the addition of sperm at t=0 min. Time in hours:minutes. Green and DIC channels are shown. Scale bars, 20 µm.

**Table S1:**
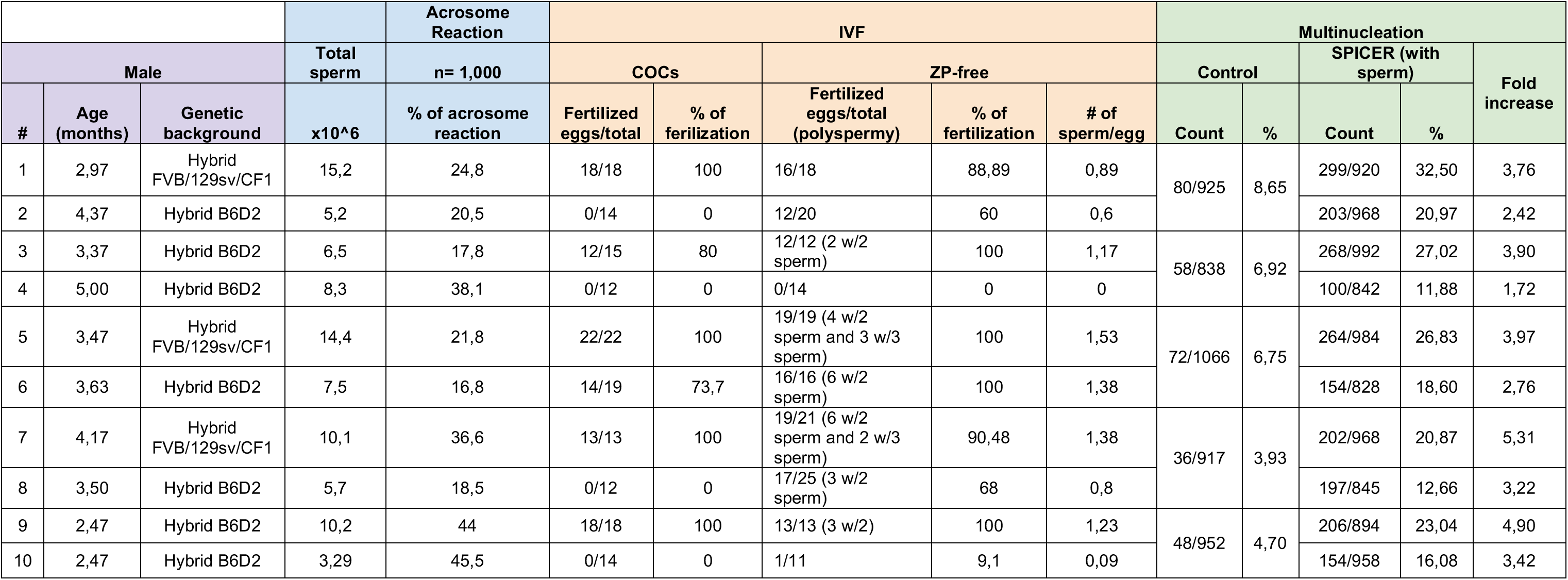
Raw data from correlation assays.

